# Newly formed place cells are stabilized by coordinated population activity during REM sleep

**DOI:** 10.1101/2022.10.10.511566

**Authors:** Richard Boyce, Hyun Choong Yong, James E Carmichael, Mark P Brandon, Sylvain Williams

## Abstract

**The involvement of rapid-eye-movement sleep (REMs) in spatial memory formation was recently demonstrated, although how neural activity during REMs influences newly-formed place field stability remains unclear. Here, we combined large-scale single-unit recordings of mouse hippocampal CA1 with an established optogenetic approach enabling REMs-selective inhibition of medial septum GABAergic neurons (MSGABA), resulting in spatial memory deficits when applied post-learning. Although individual neural activity was unaffected by REMs-selective MSGABA inhibition during a post-learning rest session, both the synchrony of population-level activity bursts observed during REMs occurring in the rest session and place field stability measured during subsequent memory recall testing were reduced vs controls. However, the latter effect was limited to place cells participating in population activity during REMs, as stability of non-participant place cells was relatively weak and indifferent between groups. This suggests that synchronous CA1 population activity during REMs stabilizes spatial representations in a plastic subpopulation of participating CA1 neurons.**

## Introduction

The collective activity of place cells, hippocampal neurons which fire in different locations of a given area ^1,2^, is thought to represent a cognitive map of a spatial environment ^3^. The long-term stability of such cognitive maps has been linked to memory performance ^4,5^, and work completed over the past several decades has shed much light on the mechanisms promoting stabilization of place cell representations. In particular, offline replay of place cell sequences observed during prior exploration of an environment has been shown to occur during highly synchronous periods of hippocampal neural activity, known as sharp-wave ripple complexes (SPW-Rs), which occur during periods of quiet (resting) wakefulness as well as non-REM sleep (NREMs) ^6,7,8^ and may play a key role in the transfer of hippocampal spatial representations to neocortical targets ^9,10^. Furthermore, the disruption of SPW-R activity following hippocampus-dependent memory encoding led to impaired memory performance ^11,12,13^, cumulatively providing strong evidence that offline SPW-R activity plays a key role in spatial memory formation.

While memory-associated SPW-R activity is largely restricted to periods of quiet wakefulness and NREMs, a direct causal relationship between REM sleep (REMs) and spatial memory formation was recently demonstrated for the first time ^14^. However, there remains a lack of mechanistic insight into precisely how offline neural activity occurring during REMs might influence the processing of spatial information. This is primarily due to an absence of experiments which have performed large-scale single unit recordings of hippocampal place cell activity under conditions in which neural activity during REMs has been manipulated with high temporal precision following encoding of spatial information. Here, we aimed to address this issue by performing microdrive tetrode recordings in area CA1 of mice expressing archaerhodopsin (ArchT) in GABAergic neurons of the medial septum (MSGABA), an established optogenetic approach ^14^ which allowed us to track the activity of many CA1 place cells across all portions of an experiment in which spatial memory deficits were induced following learning via REMs-selective silencing of MSGABA. We report that REMs-specific MSGABA inhibition did not produce any apparent alteration in activity characteristics at the level of the individual CA1 neuron during a rest session following spatial learning. However, analysis of neural activity at the population level revealed the presence of synchronous population events during REMs which were disturbed by REMs-selective MSGABA inhibition. In Control mice, the stability of spatial representations (place cell stability) of participant CA1 neurons measured in a subsequent spatial memory recall session was significantly enhanced compared to CA1 neurons which did not participate in population events during REMs; however, this enhanced stabilization was blocked by REMs-selective MSGABA inhibition during the prior rest session, suggesting that synchronous population events in CA1 during REMs contribute to the formation of spatial memory by enhancing the stability of spatial representations in a subpopulation of plastic neurons.

## Results

### Single-unit CA1 recordings in mice with ArchT expression targeted to GABAergic neurons of the medial septum

In order to achieve high temporal resolution inhibitory control over MSGABA activity, we performed intraseptal injections of a virus encoding ArchT fused to an eYFP reporter in male VGAT::Cre mice. The resulting construct (ArchT-eYFP) expression was localized to the MS-diagonal band of broca (DBB) and typically remained stable and effective for at least 4 months post-injection (Fig 1A-B). Consistent with prior reports ^14,15^, septo-hippocampal projections were found to heavily innervate the dentate gyrus, CA3, and CA1 stratum oriens regions of the hippocampus (Fig 1B, right). After allowing adequate time for full construct expression, mice were implanted with microdrives containing 4 independently movable tetrodes which were gradually lowered to the CA1 cell layer of the dorsal hippocampus over the ensuing weeks (Fig 1A-B, right) to ensure long-term stable recordings (Fig 1C). Once the lowering procedure was completed, experimental recordings commenced.

**Fig. 1.**
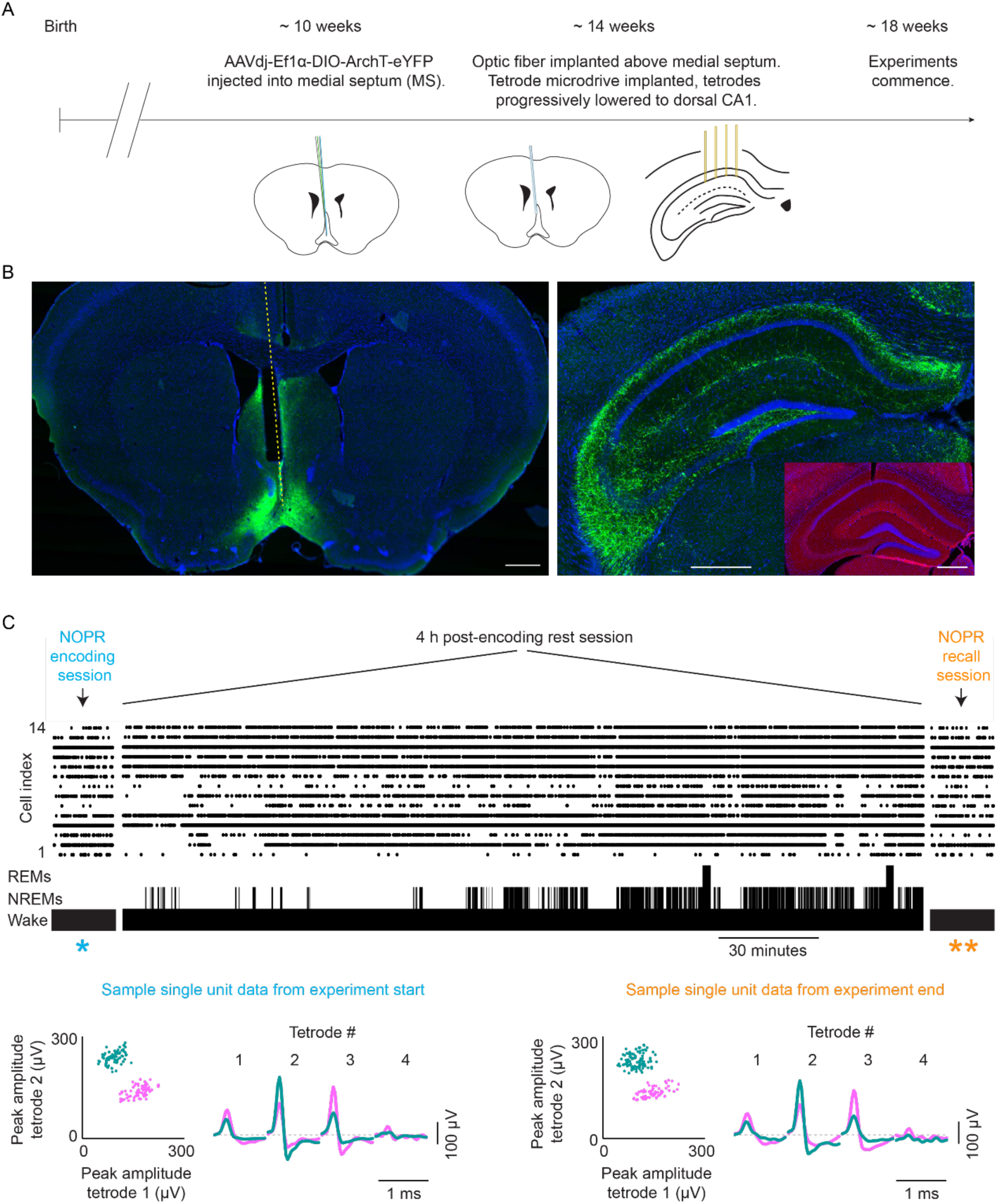
Single-unit CA1 recordings in mice with ArchT expression targeted to GABAergic neurons of the medial septum. (A) Experimental timeline. To enable selective expression of ArchT to GABAergic neurons of the medial septum (MS), an adeno-associated viral vector encoding ArchT-eYFP expression was injected into the MS of ∼10 week old VGAT::Cre male mice. ∼4 weeks later, an optic fiber was implanted with the tip placed just above the MS transfection zone to facilitate light-induced optogenetic inhibition of MS GABAergic neurons (MSGABA) below. During the same procedure, a microdrive holding 4 independently moveable tetrodes was implanted above the dorsal hippocampus. Over the ensuing ∼ 4 weeks, the tetrodes were gradually lowered into the CA1 cell layer for long-term stable recordings. (B) Left. YFP (green) and DAPI (blue) labelled coronal section taken at the level of the MS. The tract of the injection needle used to induce virus-mediated ArchT expression in MSGABA neurons is indicated by the dashed yellow line. The tract resulting from placement of the optic fiber with the tip located just above the MS is also present. Right. Dorsal hippocampal coronal section showing the presence of septo-hippocampal MSGABA projections in the dentate gyrus, CA3, and CA1 stratum oriens. Inset. Example of a tetrode tip located in the CA1 cell layer of a different DAPI-labelled dorsal hippocampal coronal section. Background color (red) has not been filtered to better resolve the full electrode tract. White scale bars in all figures = 500 µm. (C) Top. Example spike raster constructed using stable isolated dorsal CA1 neural recordings spanning all portions of a single experiment (encoding session, post-encoding rest session, recall session; encoding and recall sessions truncated for space considerations). Note that white spaces in the hypnogram represent breaks in otherwise continuous recordings when mice were transferred between recording environments. Bottom. Example of procedure for separating single neural units and ensuring long-term stability of recordings. Single units were isolated using feature (left in panel) and waveform (right in panel) plots. Only well-separated ‘clusters’ were utilized. Long-term stability was assessed by directly comparing plots compiled using data from the start (panel at left) and end of recordings (panel at right). Note that additional spikes have been removed from the plots shown for clarity.

### Optogenetic MSGABA silencing during REMs impairs novel object place recognition memory

To gain mechanistic insight into the role of hippocampal neural activity during REMs in spatial memory formation, we selectively silenced MSGABA during REMs occurring in a 4 h rest session that was interposed between the encoding and recall phases of a hippocampus-dependent spatial novel object place recognition test ^14,16^ (Fig 2A). Single unit activity was recorded for the entire duration of experiments (i.e., encoding, rest, and recall sessions) (Fig 1C) using the same recording configuration, allowing for tracking and analysis of the activity of each individual stable CA1 neuron, including that of cells with significant place fields during the encoding and recall sessions (i.e., place cells), throughout the entire experiment. At the start of an experiment mice were placed into a square open field with 2 identical objects (called ‘Object 1’ and ‘Object 2’) located in randomly assigned quadrants, and subsequently allowed 30 minutes to explore the test area. There was no difference in the amount of time mice spent exploring either object during this session (Fig 2D, left), indicating neutral preference for objects upon initial exposure. Mice were then transferred to a separate cage where they were allowed a 4 h rest period, during which sleep activity was continuously monitored via hippocampal local field potential (LFP) and electromyogram (EMG) recordings. Upon entry into REMs, MSGABA were silenced via orange light delivery to the MS transfection zone until the end of the REMs episode, at which time the light was turned off. This procedure was repeated for the entire rest session resulting in MSGABA being silenced for on average 82.6 ± 0.2 % of all REMs. While this manipulation had no effect on overall sleep activity as assessed by comparison of sleep quantity and quality to controls (Fig 2C; Fig S1B; Table S1), there was a significant reduction in REMs theta (4-10 Hz) band power (Fig 2B; Fig S1C) as reported previously ^14^. Immediately following completion of the rest session, mice were transferred back to the square open field for a 30-minute spatial memory recall test session. The test area was identical to that during the encoding session, with the lone exception being that Object 2 was moved to a new location. Considering that mice preferentially investigate novelty, the bias of a mouse for exploration of Object 2 during the recall vs encoding session (Object 2 discrimination index (DI)) is considered a metric of intact spatial memory. However, mice which had MSGABA selectively silenced during REMs (ArchT mice) showed no difference in Object 2 DI between the encoding and recall sessions (Fig 2D, right), indicating that spatial memory was disturbed in the latter. In contrast, Control mice had a significantly higher Object 2 DI during the recall session relative to the encoding session. There was no difference between groups in the total distance travelled nor in the total time spent investigating either object in the encoding and recall sessions (Fig S1A). Cumulatively, these results therefore suggest that MSGABA activity occurring selectively during REMs is critical for hippocampus-dependent spatial memory formation, consistent with our prior report ^14^.

**Fig. 2.**
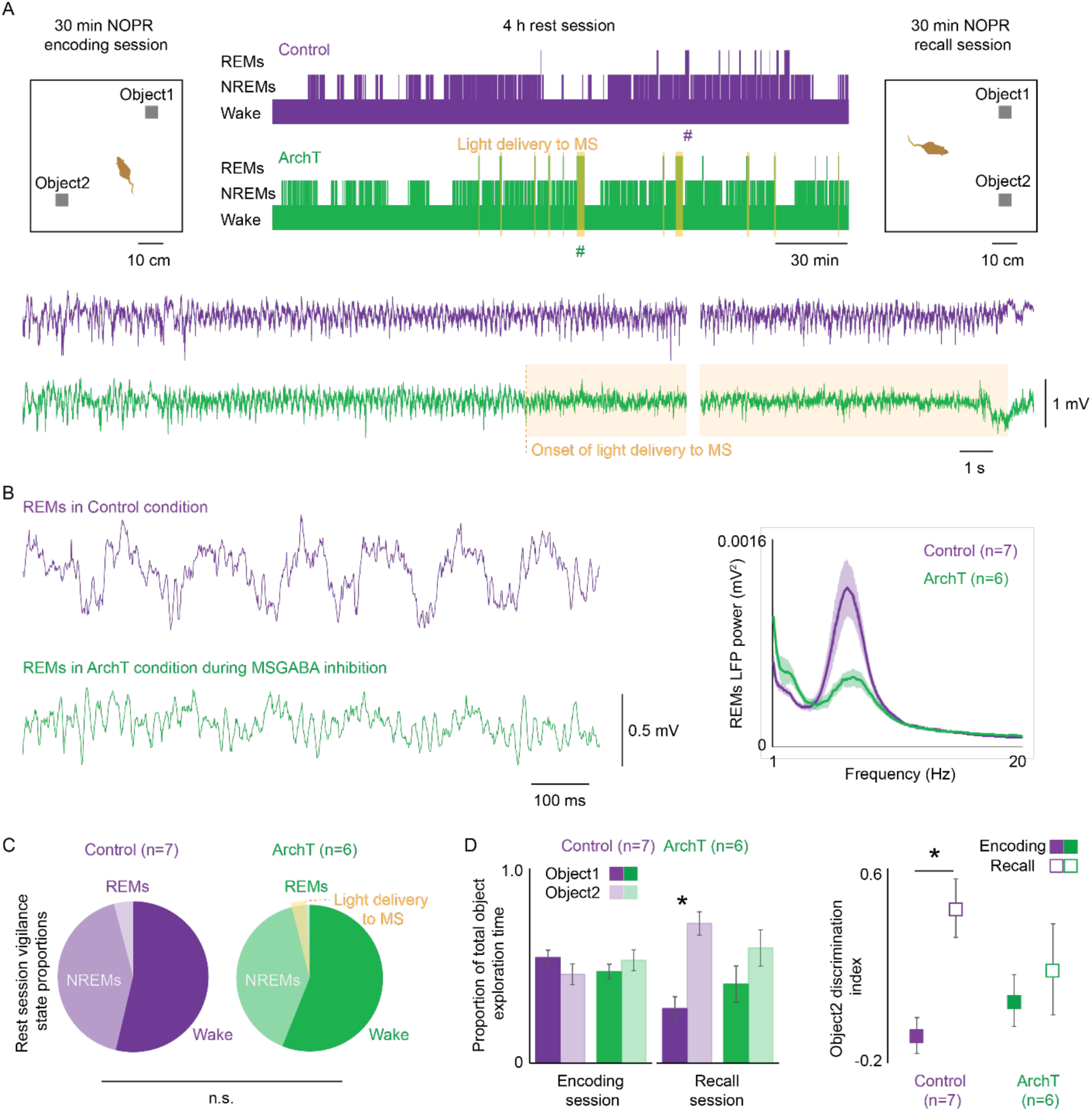
Optogenetic MSGABA silencing during REMs impairs novel object place recognition memory. (A) Schematic of behavioral experiments. Top, left. Mice were placed in a test area with two objects located in randomly assigned quadrants for a 30-minute encoding session. Top, middle. Mice were subsequently allowed a 4 h rest period during which MSGABA were optogenetically silenced selectively during REMs in the ArchT group (see sample CA1 traces below). Top, right. Following the rest period, mice were returned to the prior test area for a 30-minute spatial memory recall session. The test area was the same with the exception that one of the objects (Object 2) had been assigned to a new quadrant. (B) Left. High-resolution sample traces showing the effects of MSGABA inhibition on the CA1 LFP during REMs. Right. Spectral analysis of CA1 LFP power during REMs. Values are presented as mean ± SEM. (C) Vigilance state proportions during the rest session. Orange shading represents MSGABA inhibition. (n=7 (Control), n=6 (ArchT); n.s.=not significant, two-way ANOVA). (D) Left. Proportion of object exploration time in the encoding vs recall sessions. (n=7 (Control), n=6 (ArchT); \**P*<0.05, paired t-test). Right. Object 2 discrimination index for the encoding vs recall sessions. (n=7 (Control), n=6 (ArchT); \**P*<0.05, paired t-test). All values in D are presented as mean ± SEM.

**Fig. S1.**
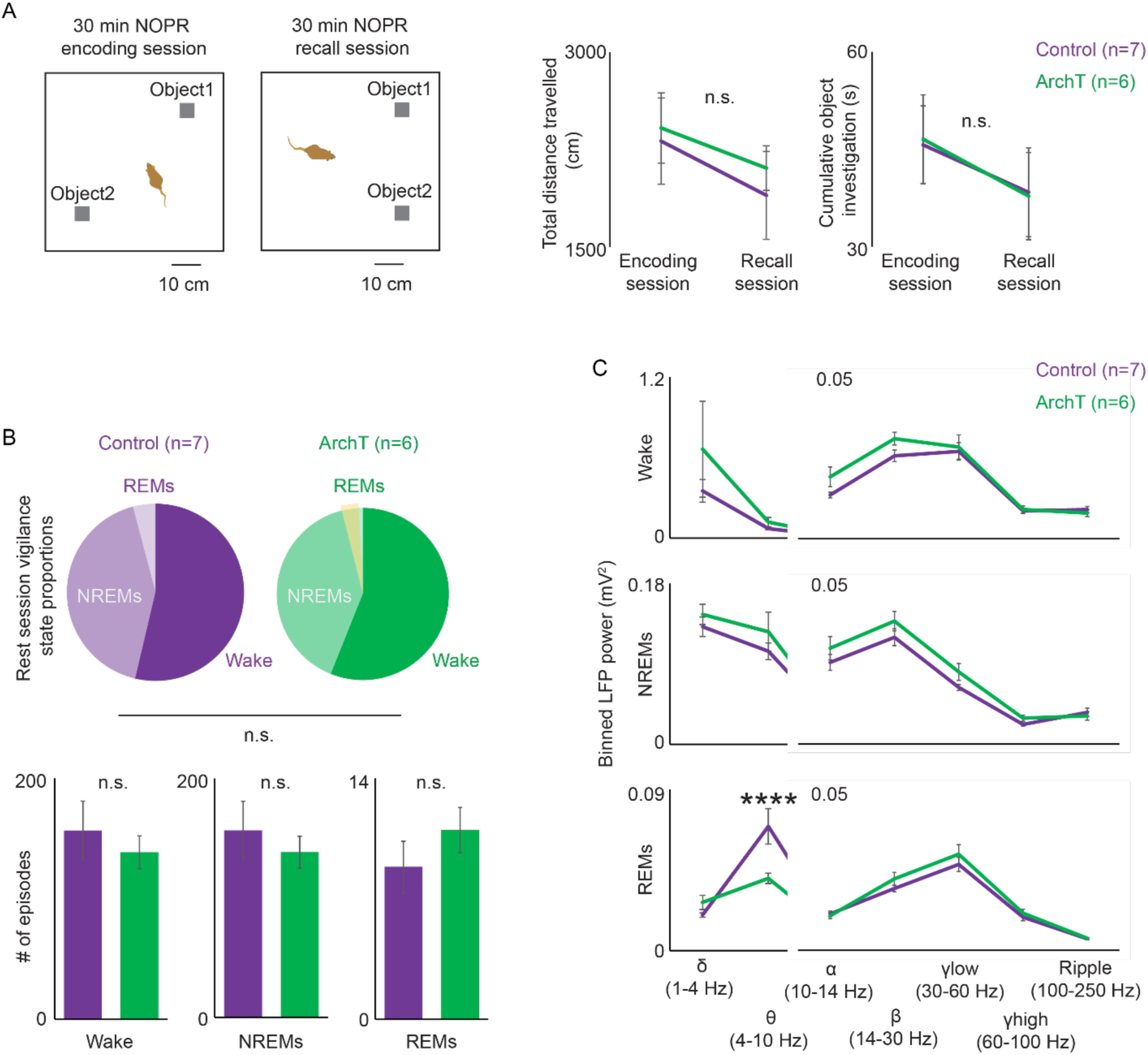
Detailed analysis of behavioral experiments. (A) Left. Test schematic of the NOPR encoding and recall sessions. Right. Quantification of the total distance travelled during the encoding and recall sessions. (n=7 (Control), n=6 (ArchT); n.s.=not significant, two-way ANOVA). Bottom, right. Total object exploration time (Object 1 exploration time + Object 2 exploration time). (n=7 (Control), n=6 (ArchT); n.s.=not significant, two-way ANOVA). Values are presented as mean ± SEM. (B) Top. Vigilance state proportions during the rest session. Orange shading represents MSGABA inhibition. (n=7 (Control), n=6 (ArchT); n.s.=not significant, two-way ANOVA). Bottom. Total number of vigilance state episodes during the rest session. (n=7 (Control), n=6 (ArchT); n.s.=not significant, two-way ANOVA). Values are presented as mean ± SEM. (C) Vigilance state-specific spectral analysis during the rest session. (n=7 (Control), n=6 (ArchT); \*\*\*\**P*<0.0001, two-way ANOVA with Šídák’s multiple comparisons test). Values are presented as mean ± SEM.

### REMs-selective MSGABA silencing has no detectable effect on CA1 neural homeostasis during the rest session

To begin our investigation into the potential neural mechanisms behind the observed disruption of spatial memory following REMs-specific MSGABA inhibition, we first focused on the activity of individual neural units isolated from tetrodes located in the CA1 cell layer during the 4 h rest session. During the rest session for mice in both the Control and ArchT groups, the first hour typically consisted primarily of periods of wakefulness with relatively sparse and brief bouts of NREMs activity, which became more consolidated and numerous from the second hour onwards (e.g., see Fig 2A). REMs was not observed in any mice during the first hour of the rest session, but began to be observed in a portion of mice during the second hour (REMs was observed in 3/7 mice (43 %) in the Control group and 2/6 (33 %) mice in the ArchT group), with all mice in both the Control and ArchT groups exhibiting multiple (at least 2) REMs episodes >60 s in duration by the fourth (final) hour of the rest session. As an initial step for our analysis, we found no difference between groups in the total number of neurons isolated during the 4 h rest period (Control (n=7) = 21.8 ± 5.5 cells, ArchT (n=6) = 20.3 ± 2.9 cells; *P*=0.83, unpaired t test). Considering past work suggesting that sleep may play an important role in the homeostatic restoration of neural firing rates to baseline levels following periods of Hebbian potentiation during prior wakefulness (the sleep homeostasis hypothesis) ^17,18,19,20,21^ along with our experimental model which involves manipulating neural (MSGABA) activity selectively during REMs, we next turned our attention to assessing whether a disruption of neural homeostasis due to REMs-selective MSGABA inhibition might provide a potential explanation for the spatial memory deficits we observed during the recall session in ArchT mice relative to Controls. As a first step, because prior work has suggested that the hippocampal theta rhythm power during REMs may serve a homeostatic role by reducing overall firing rates of hippocampal neurons while increasing ripple-associated neural synchrony observed during NREMs in a process relying on tandems of NREMs-REMs bouts ^19^, we similarly first wanted to focus on how neural firing rates evolved across several NREMs-REMs cycles in our own experiments. We therefore analyzed these features from the last continuous 30 s period of NREMs prior to the first REMs episode as well as the first continuous 30 s periods of NREMs to follow the last REMs episode of the rest session to determine whether similar processes may have been disrupted by REMs-selective MSGABA inhibition and resultant theta power attenuation (Fig 3A, top). However, we were unable to find a difference in either REMs or NREMs firing rates between the ArchT and control groups (Fig 3B, top 2 plots at right), nor in the proportion of ripple-coincident CA1 unit activity occurring during the same NREMs periods (Fig 3Ci-ii). Because the above analysis did not include periods of wakefulness and tended to primarily include NREMs data from the last portion of the rest session as this is where the majority of consolidated REMs episodes occurred, we next took a more general approach by comparing how vigilance state-specific firing rates evolved for periods of wakefulness and NREMs across the 4 h rest session (Fig 3A, bottom). For a given vigilance state, only continuous episodes of at least 30 s in duration contributed to the calculation of data values. We found no evidence of a difference in the neural firing rate distributions and mean firing rate during periods of wakefulness occurring in the first vs fourth hours of the rest session either within or between groups (Fig 3B, bottom right plot), nor in those occurring during periods of NREMs in the second vs fourth hours (neural firing rates during the first hour of NREMs were not considered for analysis due to the relatively sparse and brief nature of episodes observed during this time) (Fig 3B, plot at right, 2^nd^ from bottom). Ripple-coincident CA1 unit activity (Fig 3Ci-ii) occurring during the same NREMs periods were also indifferent between groups. For the above basic firing rate measurements, we also performed analysis of the evolution of individual neural firing rates between the different time periods sampled for each vigilance state. This was done by plotting firing rates from one time period against the other for every single cell in order to assess the possibility of firing rate redistribution amongst the entire population, finding no evidence for such an effect as data points were largely centered around the unity line (Fig 3B, plots at left). Cumulatively, our inability to find clear evidence of a difference in neural homeostasis in Control vs ArchT mice during the rest session, let alone little evidence of significant change for within-group values across the rest session, argue against the spatial memory deficits observed during the recall session in ArchT mice having been due to REMs-selective MSGABA inhibition-induced alterations in neural firing rate homeostasis.

**Fig. 3.**
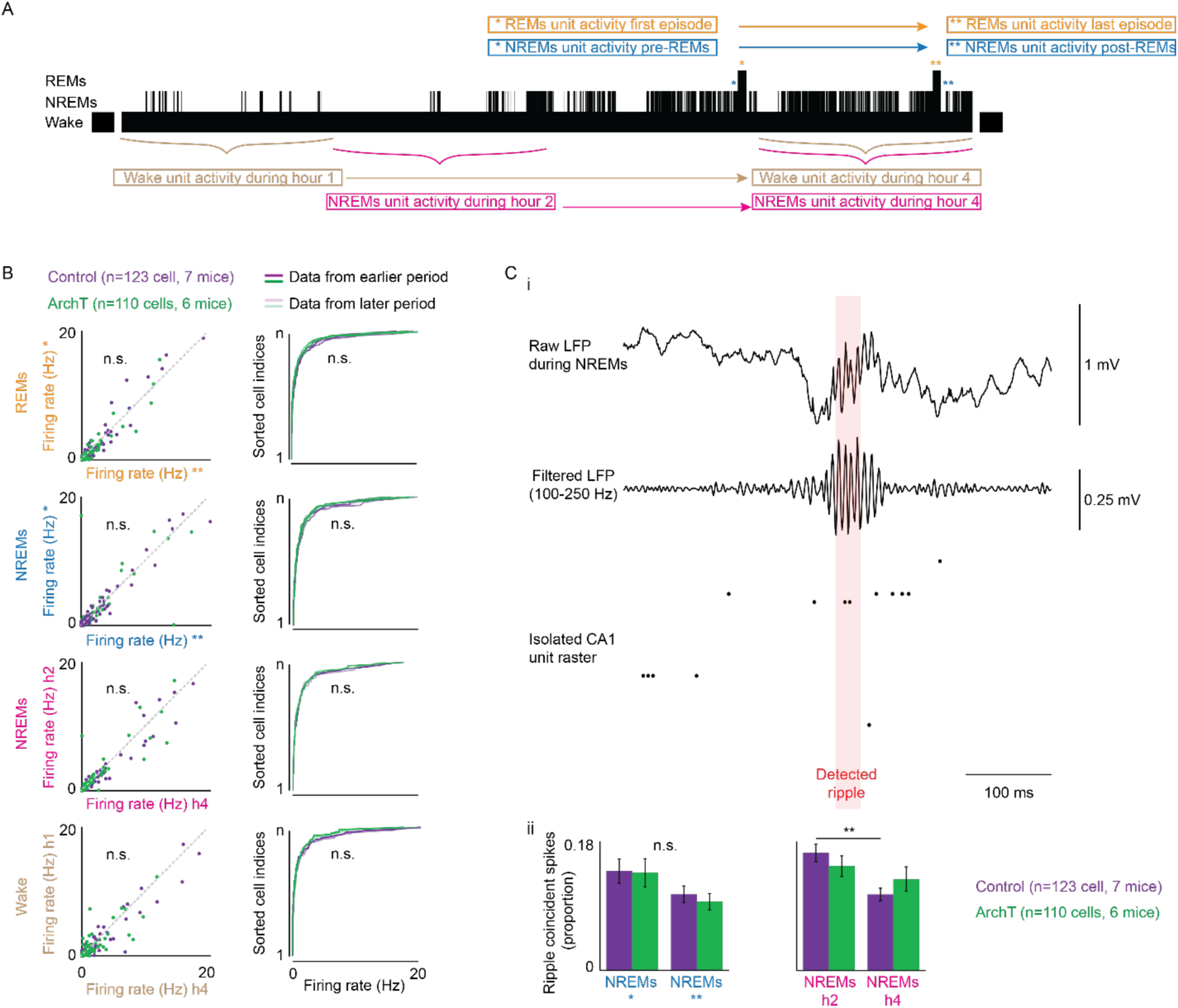
REMs-selective MSGABA silencing has no detectable effect on CA1 neural homeostasis during the rest session. (A) Example hypnogram from a rest session showing the approximate locations of vigilance state-specific time points used for subsequent analysis. (B) Left. Pooled state-specific firing rate plots of CA1 single units from Control vs ArchT mice showing the evolution of firing rates measured at different timepoints of the rest session (indicated by labelling and depicted in the accompanying hypnogram). Dashed grey line indicates unity. (n=123 cells from 7 mice (Control), n=110 cells from 6 mice (ArchT); n.s.=not significant, Mann Whitney test of mean change in firing rate for each cell at different timepoints). Right. Pooled state-specific firing rate plots of CA1 single units from Control vs ArchT mice. Data were sorted in ascending firing rate prior to plotting and plots are scaled according to the total n to allow for visual comparison between firing rate distributions between Control and ArchT conditions. (n=123 cells from 7 mice (Control), n=110 cells from 6 mice (ArchT); n.s.=not significant, Mann Whitney test of average firing rate values). (C,i) Example of ripple coincidence analysis of CA1 single unit activity during NREMs. Significant ripple activity was detected from filtered traces using a thresholding approach (see method details). For each timepoint analyzed, the proportion of spikes from each isolated CA1 unit which coincided with ripples detected within the analysis period (red shading) was quantified and values from all cells were pooled. (ii, left) Plot of ripple-spike coincidence values calculated for pre- vs post-REMs periods of NREMs (n=123 cells from 7 mice (Control), n=110 cells from 6 mice (ArchT); n.s.=not significant, Mann Whitney test). (ii, right) Plot of ripple-spike coincidence values calculated for NREMs occurring within the 2^nd^ vs 4^th^ hours of the rest session (n=123 cells from 7 mice (Control), n=110 cells from 6 mice (ArchT); \*\**P*<0.01, Mann Whitney test, Control hour 2 vs hour 4 of rest session). Values in Cii are presented as mean ± SEM.

### REMs-selective MSGABA silencing has no detectable effect on phase-locking of CA1 neural activity to theta oscillations during REMs

During REMs in rodents, a key feature of hippocampal LFP activity is the ∼7 Hz theta rhythm (e.g., Fig 2A-B). Although relatively scarce, prior reports have presented evidence that the activity of individual CA1 neurons occurring during REMs following completion of a behavioral task is locked to specific phases of the theta-rhythm ^22,23^, perhaps as a function of the recency of place field formation for a given neuron ^22^ (but see ^23^). As previous work has suggested that the phase of the theta oscillation at which a CA1 neuron fires plays an important role in long-term potentiation (LTP), with activity at the peak versus trough being associated with LTP and long-term depression (LTD), respectively ^24,25,26,27^, the observation of such phase-locking activity during REMs has been suggested to influence the processing of spatial information ^22^. We therefore performed a similar analysis here to determine whether a general disturbance in phase-locking of isolated neural activity to theta oscillations during REMs in the ArchT group due to theta rhythm power attenuation could be behind the spatial memory deficits subsequently observed during the recall session (Fig 4A-B). However, phase-locking of CA1 unit activity to theta oscillations was negligible in both the Control and ArchT groups (Fig 4C, top). To ensure that this result was not due to the inclusion of data from cells which did not form significant spatial representations while performing the spatial memory test (i.e., non-place cells), we also performed the above analysis using data from significant place cells only, obtaining a similar result (Fig 4C, bottom). These results argue against disruption of the preferred theta rhythm firing phase of individual neurons during REMs as a cause of the subsequently observed spatial memory deficits.

**Fig. 4.**
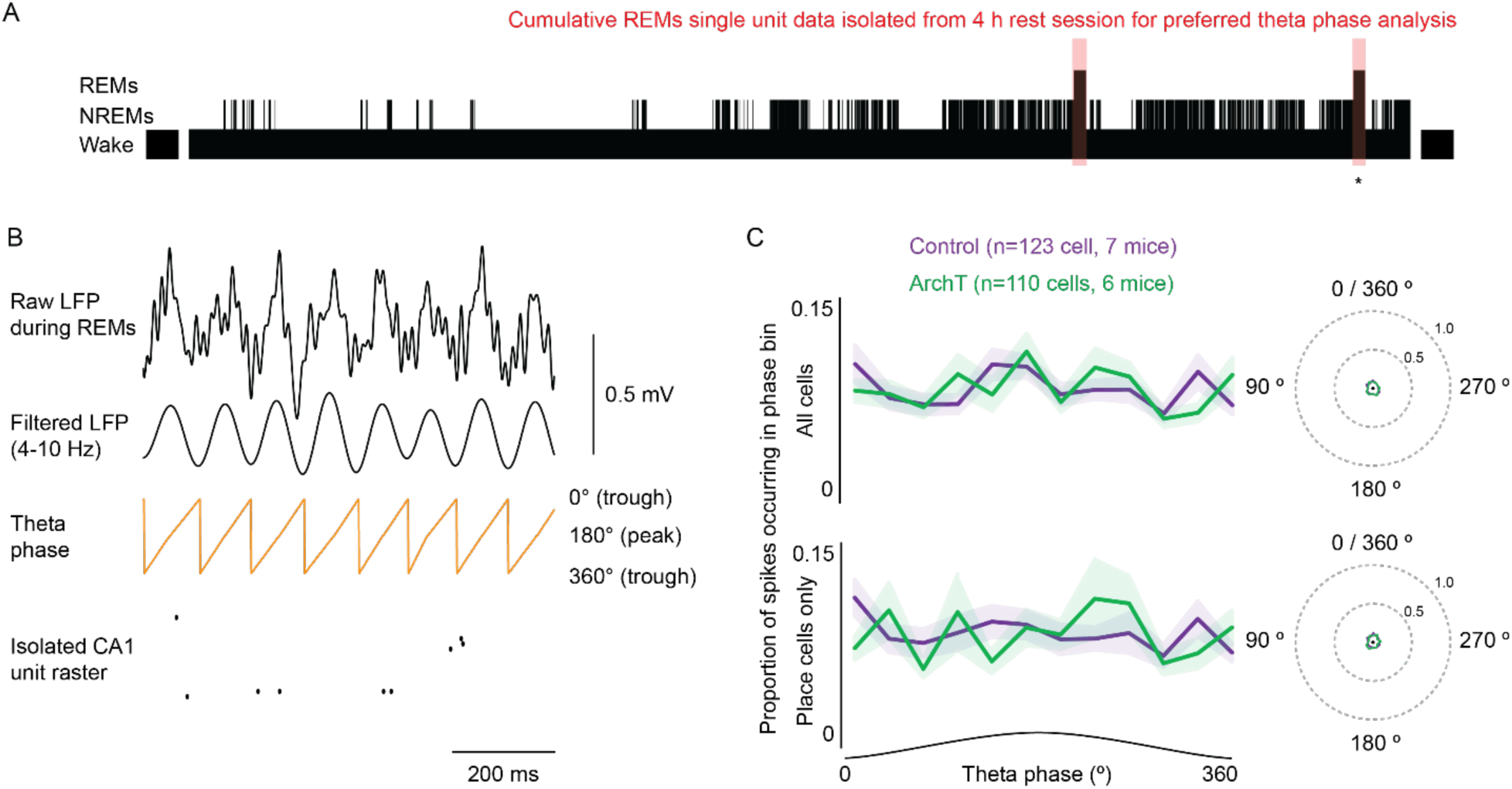
REMs-selective MSGABA silencing has no detectable effect on phase-locking of CA1 neural activity to theta oscillations during REMs. (A) Example hypnogram from a rest session depicting selection of cumulative REMs data for theta phase-locking analysis of isolated neural activity. (B) Example of theta phase calculation during REMs. Raw LFP traces were first bandpass filtered at theta (4-10 Hz) frequency and the theta phase at each timepoint was subsequently found (orange trace), enabling the firing phase of each spike from every isolated neuron to be calculated. Asterisk indicates approximate location of trace in experiment. (C) Analysis of the relationship between CA1 single unit activity and theta phase during REMs for all neurons (top) as well as neurons which had significant place fields calculated during spatial memory testing (i.e., place cells, bottom). Extent of shading indicates standard error. At right, radial plots of preferred phase distributions overlay mean vectors, which are not visible due to negligible values.

### REMs-selective silencing of MSGABA reduces neural synchrony during CA1 population event activity

Although the majority of REMs was characterized by generally sparse and uncoordinated CA1 neural activity with the exception of a relatively small number of neurons which occasionally exhibited tonic activity, we consistently observed the presence of transient synchronous CA1 population activity ∼10 ms in duration (Fig 5A-B). Thus, while we were unable to find differences in the characteristics of isolated CA1 neural unit activity between groups, we next hypothesized that synchronous CA1 population activity during REMs could have a role in spatial memory formation which might have been disrupted in ArchT mice. To evaluate this possibility, we identified synchronous population events automatically by using a windowing-based approach to determine 10 ms segments of REMs with atypically high (above significance threshold) levels of neural coactivity (Fig S2A-B, C (left); also see method details). The relative number of significant synchronous population events occurring during REMs was reduced in the ArchT group versus Controls, although this difference was not significant (Fig S2C, right). However, when we analyzed synchrony of neural activity occurring during identified population event activations by determining the minimum interval of time within the corresponding 10 ms window during which the number of active CA1 neurons (neurons firing at least 1 spike) was equal to or greater than the experiment-specific coactivity threshold for each activation (method details), we found that average values were significantly higher in the ArchT group relative to controls (Fig 5B-C). In particular, there was a clear lack of the most highly synchronous events occurring during the control conditions (<8 ms interval between first and last spike of a given population event) in experiments of the ArchT group. As a final step, we wanted to investigate the characteristics of LFP activity occurring at the time of population events to see if we could identify specific factors associated with the disrupted synchrony of population events during REMs in the ArchT group experiments. Unlike SPW-Rs occurring during NREMs, there were no overt oscillatory features associated with REMs population events in the LFP signal. Wavelet power was thus calculated for 10 ms time windows centered around population event time points (defined as the mid-point of the temporal bounds of a given population event (method details)) and averaged for each condition. Both ArchT and Control mice had activity peaks in the ∼60-100 Hz ‘high gamma’ range that was preferentially coupled to the theta trough (Fig 5D, i,ii). However, in addition to the characteristic reduction in theta power in ArchT mice, wavelet analysis further revealed the presence of prominent theta peak-associated activity in the ∼30-60 Hz ‘low gamma’ oscillation range during REMs population events in these mice that was comparatively small in Controls, perhaps due to increased influence of intrahippocampal vs extrahippocampal gamma rhythm generators resulting from MSGABA inhibition ^28,29^. This may have promoted reduced synchrony of population events relative to weaker oscillatory activity in the 60-100 Hz range that was present in both ArchT and Control mice, as phase-locking of population event activity to ∼30-60 Hz and ∼60-100 Hz oscillatory activity was relatively strong and preferentially occurred at the peak in both Control and ArchT mice (Fig. 5D, iii.). The latter results were not likely to be due to spike contamination of LFP signals ^30^, as the preferred theta phase of neural population events and high gamma oscillations (theta phase of maximal high gamma power) showed opposite trends.

**Fig. 5.**
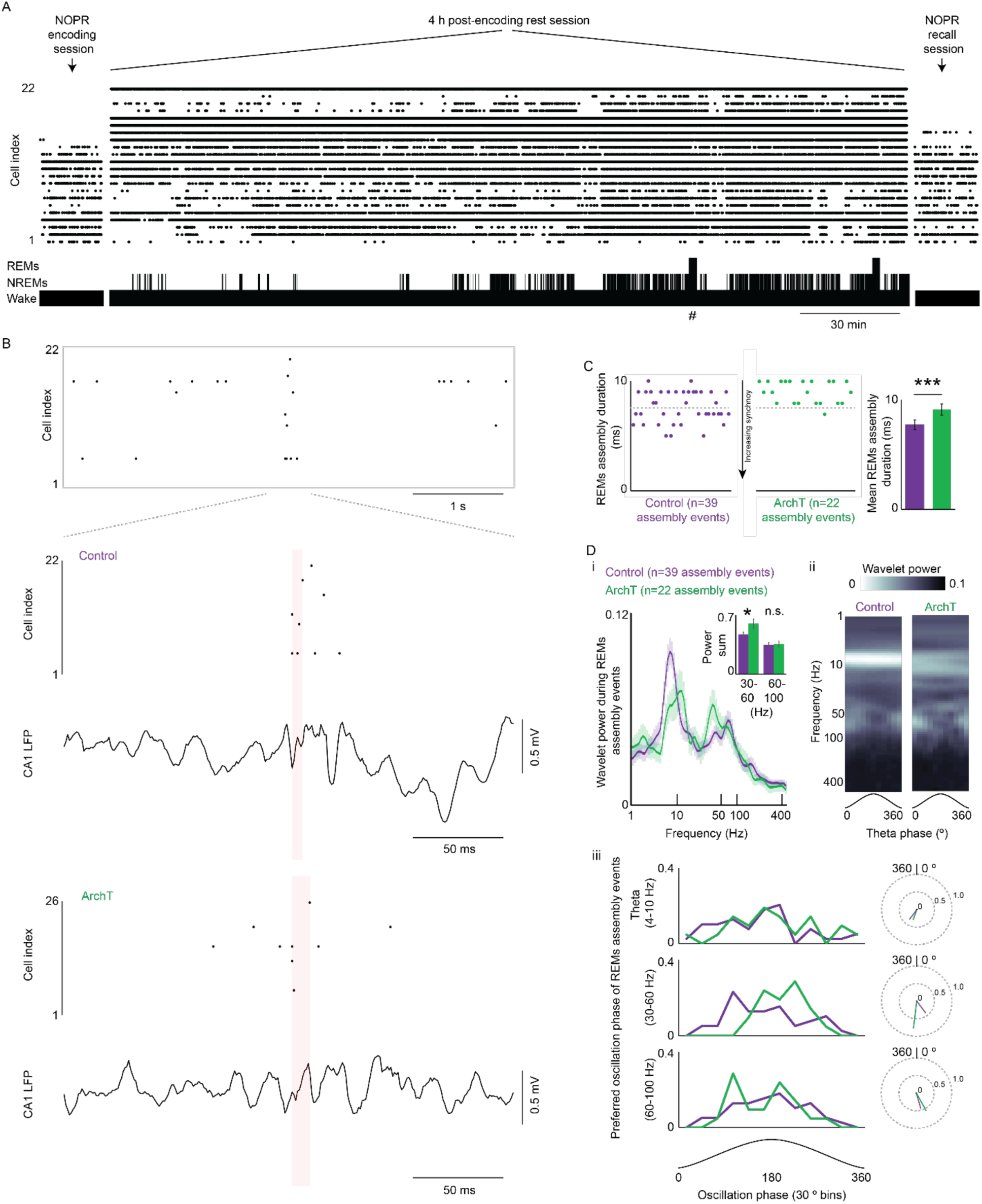
REMs-selective silencing of MSGABA reduces neural synchrony during CA1 population event activity. (A) Example isolated CA1 unit recording from a single Control experiment. The activity of each individual single unit was tracked across the entire duration of the experiment (encoding, rest, and recall sessions). Neurons have been organized such that neurons active during encoding and/or recall sessions, as well as the rest session, are plotted first. (B) Top. Spike raster centered around an example detected population event occurring during the experiment shown in (A). The # symbol in (A) indicates the approximate location of the example detected population event. Middle. High-resolution view of the detected population event with the corresponding LFP trace below. The location and calculated duration (see method details) of the sample detected population event is indicated by the pink shading. Bottom. Same as middle, except showing example data from a paired ArchT experiment. (C) Pooled analysis of detected REMs population event durations during the rest session. (n=39 detected population events from 7 mice (Control), n=22 detected population events from 6 mice (ArchT); \*\*\**P*<0.001, Mann Whitney test). Plot at right is presented as mean ± SEM. (D) Analysis of LFP characteristics during REMs population events. (i) Pooled wavelet power spectra calculated for a 10 ms window centered over REMs population events during the rest session. Inset. Binned wavelet power for ‘low gamma’ (30-60 Hz) and ‘high gamma’ (60-100 Hz) frequency bands. (n=39 detected population events from 7 mice (Control), n=22 detected population events from 6 mice (ArchT);\**P*<0.05, n=two-way ANOVA with Šídák’s multiple comparisons test. Values are presented as mean ± SEM. (ii) Average theta phase-amplitude coupling plots for Control vs. ArchT conditions, constructed by pooling phase-amplitude coupling data from the theta cycle coinciding with each detected REMs population event. (iii) Pooled analysis of the preferred theta (4-10 Hz), ‘low gamma’ (30-60 Hz), and ‘high gamma’ (60-100 Hz) frequency oscillation phase of detected population event activity during REMs. The time point of population event activity was taken as the mid-point of the activation window. The corresponding mean vector length calculations are shown at right.

**Fig. S2.**
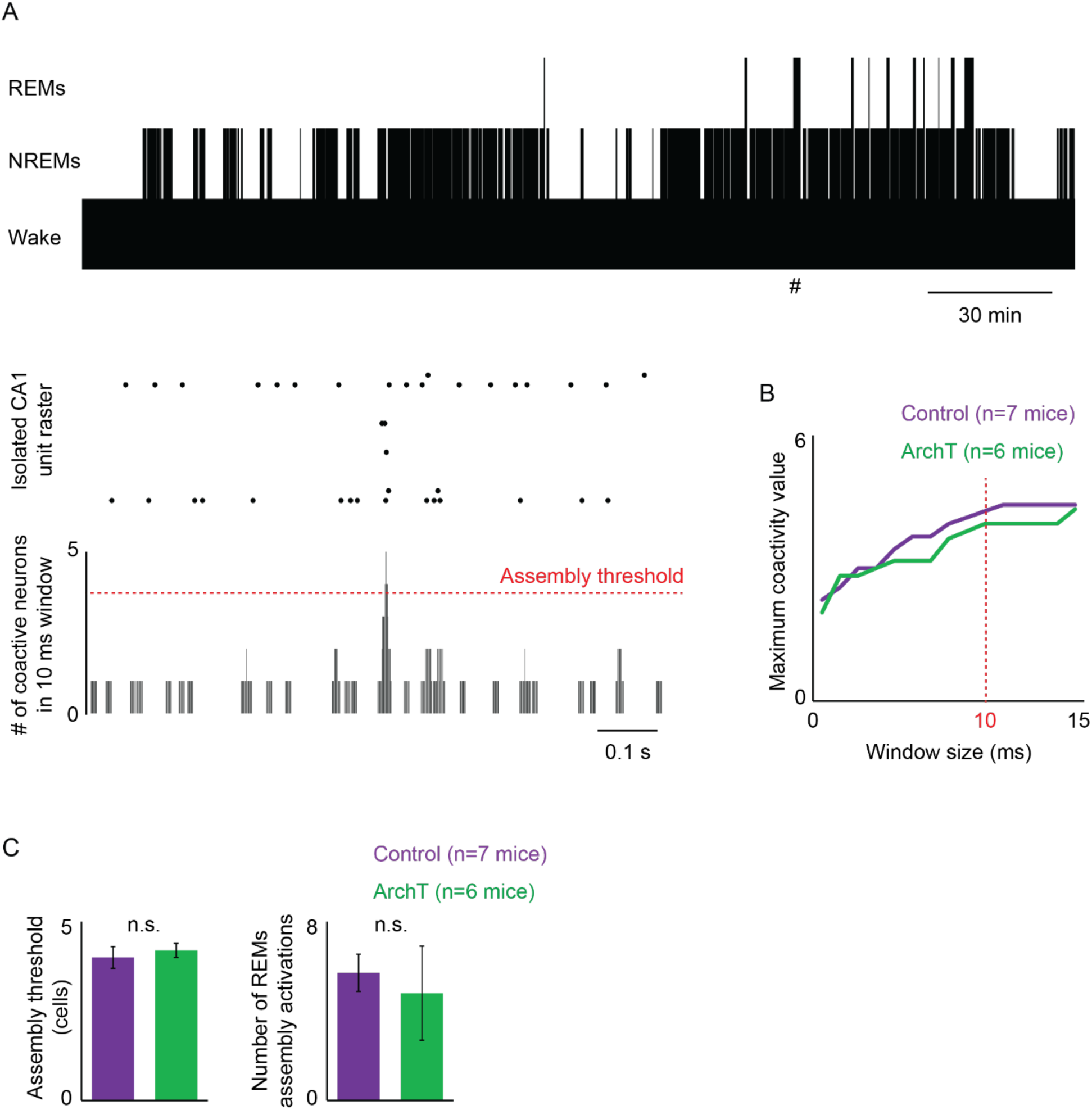
Detection of CA1 population events during REMs. (A) Top. Example hypnogram from a rest session showing the approximate location of REMs CA1 spike raster data shown below. Middle. CA1 spike raster data during REMs centered around a detected population event. Bottom. Population event activity was detected by sliding a 10 ms window along each data point of REMs and calculating the total number of neurons which fire at least 1 spike within the window for each. Data points corresponding to spike coactivity values higher than the calculated experiment-specific threshold (mean + 5 standard deviations of the coactivity value for all REMs data points rounded up to the nearest whole number; see method details) were considered to be locations of population event activity. Significant values detected at consecutive data points were counted as the same population event. (B) Maximum coactivity values calculated during REMs using a windowing approach with different window sizes. A window size of 10 ms was selected for use in all subsequent analyses because it provided the maximum trade-off between temporal resolution and the ability to capture CA1 spike coactivity, as indicated by the leveling off of the curve at ∼10 ms. (C) Characteristics of population event activity detected during REMs. Left. Population event threshold. Right. Quantification of population event activity during REMs. (n=7 (Control), n=6 (ArchT); n.s.=not significant, unpaired t test). Values are presented as mean ± SEM.

### MSGABA silencing during REMs reduces place cell stability in a subset of CA1 neurons

Having analyzed the characteristics of CA1 unit activity during the 4 h rest period, we next turned our attention to the basic characteristics of CA1 place cells during the encoding and recall sessions. We found no difference between groups in the number of significant place cells identified during both the encoding or recall test sessions (Control (n=7) = 12.7 ± 2.3 cells, ArchT (n=6) = 11.2 ± 2.3 cells; *P*=0.66, unpaired t test). Although not all single units identified during the 4 h rest session were active during the encoding and recall session (Control (n=7) = 22 ± 12 % single units were active during rest but not encoding+recall sessions, ArchT (n=6) = 21 ± 15 % single units were active during rest but not encoding+recall sessions; *P*=0.93, Kruskall-Wallis test), the majority of aforementioned identified place cells were also active during the intervening rest session (e.g., see Fig 6A). There were no consistent differences between groups in the mean values for key place cell characteristics (place field size, mean firing rate, maximum firing rate, information content, sparsity, and selectivity) for either the encoding or recall session (Table S2). Place cell stability between the encoding and recall sessions, assessed through analysis of place field overlap (Control (n=37 place cells from 7 mice) = 0.42 ± 0.06, ArchT (n=41 place cells from 6 mice) = 0.32 ± 0.05; *P*=0.19, Mann Whitney test) and centroid displacement (Control (n=37 place cells from 7 mice) = 17.4 ± 2.8, ArchT (n=41 place cells from 6 mice) = 21.3 ± 2.3; *P*=0.12, Mann Whitney test), was also not found to differ between any of the groups (method details). Although we were unable to detect any significant difference in general place cell characteristics between groups that could explain our observed spatial memory deficit in ArchT mice relative to controls, we next chose to concentrate on the subset of CA1 neurons which were active during REMs population events (18/37, 14/41 neurons used for stability analysis for Control and ArchT conditions, respectively (method details)). Specifically, we hypothesized that the relative reduction in population event synchrony in ArchT mice relative to Controls may have resulted in deficits in constituent place cell function during the recall session which were obscured when analyzing all neurons together. To this end, we first calculated the basic properties of REMs population event-active isolated CA1 neurons during the encoding and recall session. We found that there was no consistent difference in these measures between groups (Table S3). However, the place field centroid shift was increased, and overlap proportion decreased, on average in REMs population event-active place cells between the encoding and recall sessions in ArchT mice relative to Controls, indicating enhanced place cell stability in the latter (Fig 6A-C). This deficit was only found for REMs population event-active CA1 neurons, as analysis of REMs population event non-active CA1 neurons revealed relatively reduced place cell stability that was not different between groups. To determine whether the above results may manifest as a disruption of place cell activity at the population level, we performed a population vector analysis ^13,31,32^ of place cells by measuring correlation coefficient values obtained between population rate vectors calculated during the encoding and recall sessions on a pixel-by-pixel basis (Fig 6D). We found that the median correlation coefficient value obtained for population vector analysis between the encoding and recall sessions was significantly smaller in ArchT mice relative to controls, indicating a disruption in spatial representation at the population level in CA1 of mice which had MSGABA selectively silenced during prior REMs.

**Fig. 6.**
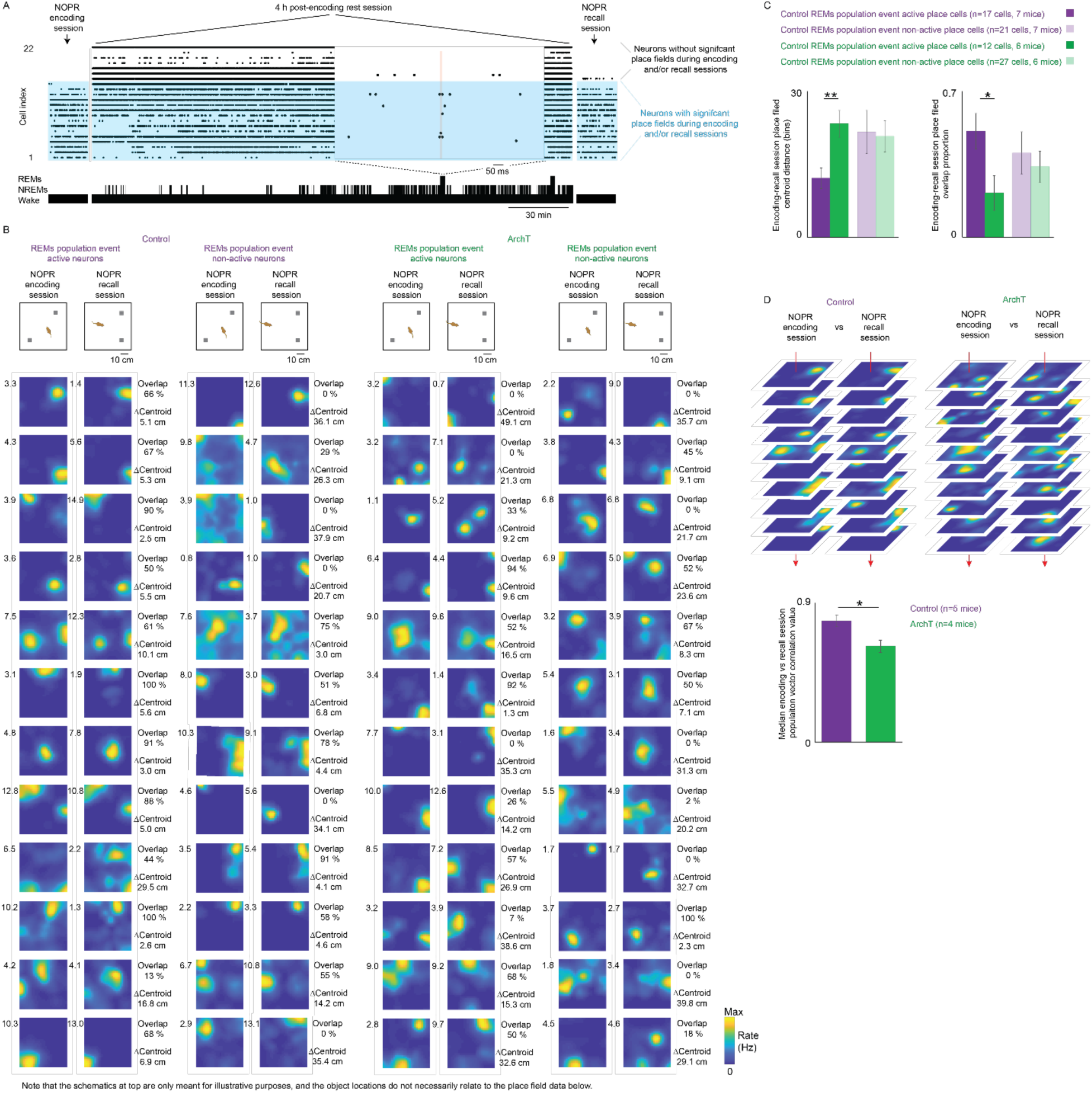
MSGABA silencing during REMs reduces place cell stability in a subset of CA1 neurons. (A) Experiment schematic demonstrating how ‘REMs population event active’ and ‘REMs population event non-active’ place cells were identified for stability analysis. Inset. High-resolution spike data (500 ms duration) centered around a single population event activation identified during REMs. The red shading indicates the location of an identified REMs population event; neurons active during at least one of these events in a given experiment were identified as ‘REMs population event active’. (B) Example of CA1 place fields calculated for the encoding and recall sessions. Values used to assess the stability of place fields across the two sessions (place field overlap and centroid displacement (see method details)) are included for each place cell at right. Max firing rates are indicated for each cell during the encoding and recall sessions by the value at top right of each corresponding rate map. Neurons which contributed to at least 1 population event detected during prior REMs have been separated from those which did not (REMs population event active and REMs population event non-active neurons, respectively). Note that the schematics at top are only meant for illustrative purposes, and the object locations do not necessarily relate to the place field data below. (C) Left. Pooled place cell encoding-recall session overlap proportions. (n=17,21 cells from 7 mice (Control REMs active and Control REMs non-active conditions, respectively), n=12, 27 cells from 6 mice (ArchT REMs active and ArchT REMs non-active conditions, respectively); \**P*<0.05, Mann Whitney test). Right. Pooled encoding-recall session centroid distance. (n=17, 21 cells from 7 mice (Control REMs active and Control REMs non-active conditions, respectively), n=12, 27 cells from 6 mice (ArchT REMs active and ArchT REMs non-active conditions, respectively); \**P*<0.05, \*\**P*<0.01, Mann Whitney test). Values are presented as mean ± SEM. (D) Place cell population vector analysis. Top. Schematic of the analysis procedure using sample data from an experiment for the Control and ArchT conditions. Place fields calculated for the encoding and recall sessions from each mouse were stacked on top of each other. Population vectors, rate values obtained from each place field map at a given pixel, were then constructed for the encoding and recall sessions and directly compared via Pearson correlation. Only experiments with data available for at least 5 significant place cells were used. Bottom. Plot of the median correlation coefficient values obtained from this analysis. (n=5 (Control), n=4 (ArchT); \**P*<0.05, Mann Whitney test). Values are presented as mean ± SEM across experiments.

## Discussion

Despite decades of research which has indicated a role for REMs in the formation of spatial memory, there is currently a lack of direct mechanistic insight into precisely how neural activity occurring during REMs might influence the processing of spatial information. Here, we combined large-scale CA1 neural unit recordings with an established optogenetic method which induced spatial memory deficits via silencing of MSGABA neurons selectively during REMs occurring during a 4 h rest session interposed between the spatial memory encoding and recall sessions. Analysis at the level of the individual CA1 neuron during the rest session did not reveal evidence of an effect of MSGABA inhibition on activity characteristics (e.g., no disruption of firing rate homeostasis or phase-locking of neural activity to prominent LFP rhythms during REMs). However, population-level analysis of CA1 neural unit activity during REMs revealed the presence of CA1 population events ∼10 ms in duration, a timescale residing within the effective window for synaptic plasticity ^33,34^, whose synchrony was disturbed by REMs-selective MSGABA inhibition. This was potentially due to the unique presence of prominent 30-60 Hz ‘low gamma’ frequency oscillatory activity, to which population event activity was phase-locked, in ArchT mice. Such activity may have promoted reduced synchrony of population event activity relative to weaker oscillatory activity in the 60-100 Hz ‘high gamma’ range that was present in both ArchT and Control mice. The precise mechanistic relation between MSGABA disruption and increased ‘low gamma’ activity is currently unclear, but could be due to a resultant increase in the influence of intrahippocampal vs extrahippocampal gamma rhythm generators ^28,29^. Subsequent analysis of REMs population event-active neurons during spatial memory recall testing revealed that they comprised a subpopulation of place cells with heightened stability relative to REMs population event non-active neurons, with the heightened stability found for place cells belonging to the former category being completely nullified by prior REMs-selective MSGABA inhibition. Cumulatively, these effects manifested as a disruption in population-level place cell stability during the recall test session relative to the encoding session.

In our study, we found no clear indication of sleep-dependent neural homeostasis in either the ArchT or Control groups, arguing against disturbed neural homeostasis via REMs-selective MSGABA inhibition being responsible for the spatial memory deficits we subsequently observed in the spatial memory recall session. Although this result is contrary to the sleep homeostasis hypothesis ^17,18,19,20,21^, previous work has also failed to produce supporting evidence (e.g.,^14,35^). The reason for such inconsistency is not known, but in the present study could be due to methodological differences such as the use of different species or obfuscation / blocking of homeostatic mechanisms by additional neural dynamics induced by the prior spatial memory encoding session ^36^.

The theta rhythm is a key feature of the LFP during REMs in rodents such as mice and rats. While a relatively sparse collection of studies has previously suggested that during REMs the activity of CA1 neurons which formed place fields in a test area prior to sleep is phase-locked to the REMs theta rhythm ^22,23^, we were unable to find any indication of consistent significant phase locking of neural activity to the REMs theta rhythm in either the ArchT or Control groups during REMs. One possible reason for this discrepancy is the use of different species, as both prior works utilized rat models. An additional difference between at least one of these studies ^22^ and our current work is that REMs used for analysis was reportedly taken from the hour immediately following the behavioral task, whereas in our experiments, REMs tended to not occur until 3 or 4 hours following the behavioral task, a difference likely due to mice being relatively unhabituated to our test area and procedure prior to recording. Thus, it cannot be ruled out here that species-specific differences or a time-dependent effect could be a factor in our inability to replicate these prior works.

In conclusion, our data has provided mechanistic insight by suggesting that synchronous neural population events occurring in CA1 during ‘offline’ periods of REMs play a key role in spatial memory by promoting enhanced stability of the spatial representations of a subset of plastic neurons following encoding of spatial information. From a broader perspective, the relatively infrequent and transient nature of isolated CA1 neural population event activity and general diversity in overt population-level CA1 neural activity patterns that we observed during REMs is consistent with it being a dynamic and heterogeneous behavioral state that is likely capable of performing multiple ^37,38,39,40,41^ and perhaps at times opposing (e.g., ^14,42^) situation-specific functions.

## Acknowledgements

RB was supported by The International Human Frontier Science Program Organisation (HFSP) postdoctoral fellowship #LT000835/2018-L while completing this work. JEC was supported by an Alzheimer’s Association Grant AARF-22-928742. SW was supported by Canadian Institutes for Health Research (CIHR) Foundation Program FDN-148478, Natural Sciences and Engineering Research Council of Canada (NSERC) Discovery Grant RGPIN-2020-06717, and a Tier 1 Canada Research Chair.

## Author contributions

RB and SW designed the experiments. RB performed virus injections and RB and HYC performed surgical implantation of the optic fiber and microdrive. RB performed all analysis and wrote the paper. RB, JEC, and SW discussed the results. MPB and SW established the laboratories in which experiments took place. SW supervised the project and provided financial support.

## Tables

**Table S1.**
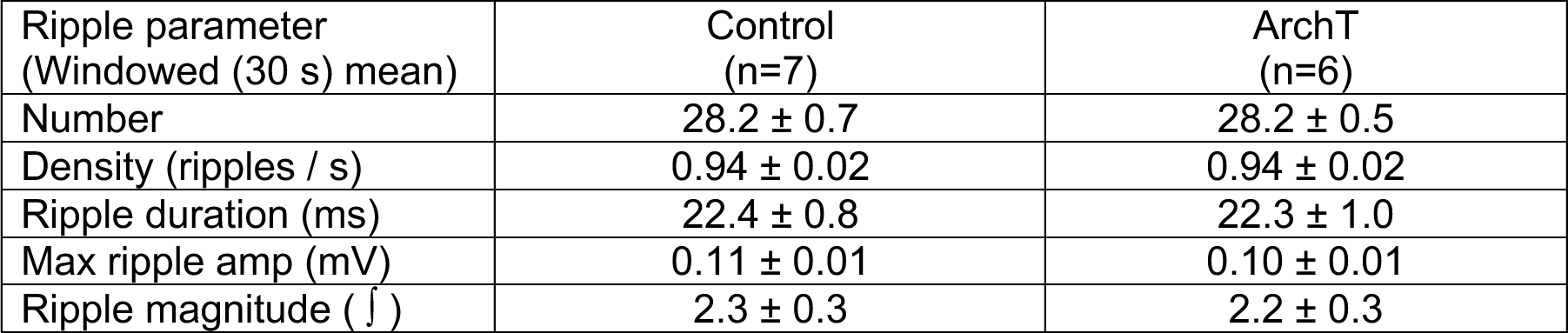
Analysis of CA1 ripple characteristics during NREMs occurring within the 4 h rest period between the encoding and recall test sessions. (n=7 (Control), n=6 (ArchT); non-significance indicated by absence of symbols, two-way ANOVA with Šídák’s multiple comparisons test). All values are presented as mean ± SEM.

**Table S2.** Analysis of place cell characteristics between Control and ArchT mice during the encoding and recall test sessions. (n=38 cells from 7 mice (Control), n=41 cells from 6 mice (ArchT); non-significance indicated by absence of symbols, unpaired t test (place field size) and Mann Whitney tests (max rate, mean rate, information content, sparsity, selectivity) were used). All values are presented as mean ± SEM.

|  |  | Size<br>(cm<sup>2</sup>) | Max rate<br>(Hz) | Mean<br>rate (Hz) | Information<br>content | Sparsity | Selectivity |
| --- | --- | --- | --- | --- | --- | --- | --- |
| Encoding<br>session | Control | 203 ± 20 | 5.0 ± 0.6 | 3.9 ± 0.4 | 1.3 ± 0.2 | 0.36 ± 0.04 | 9.7 ± 1.6 |
|  | ArchT | 175 ± 17 | 3.9 ± 0.5 | 3.1 ± 0.4 | 1.5 ± 0.1 | 0.29 ± 0.03 | 11.0 ± 1.0 |
| Recall<br>session | Control | 182 ± 18 | 4.4 ± 0.7 | 3.5 ± 0.5 | 1.6 ± 0.2 | 0.31 ± 0.03 | 11.0 ± 1.2 |
|  | ArchT | 185 ± 16 | 4.1 ± 0.5 | 3.2 ± 0.4 | 1.5 ± 0.1 | 0.28 ± 0.02 | 11.6 ± 1.7 |

**Table S3.**
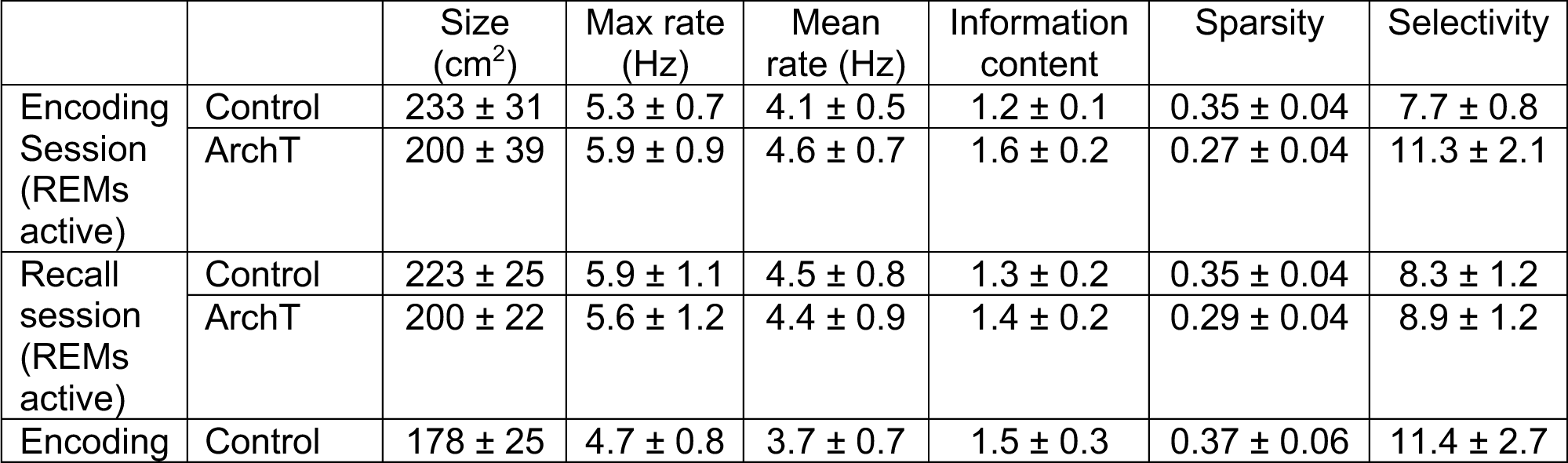
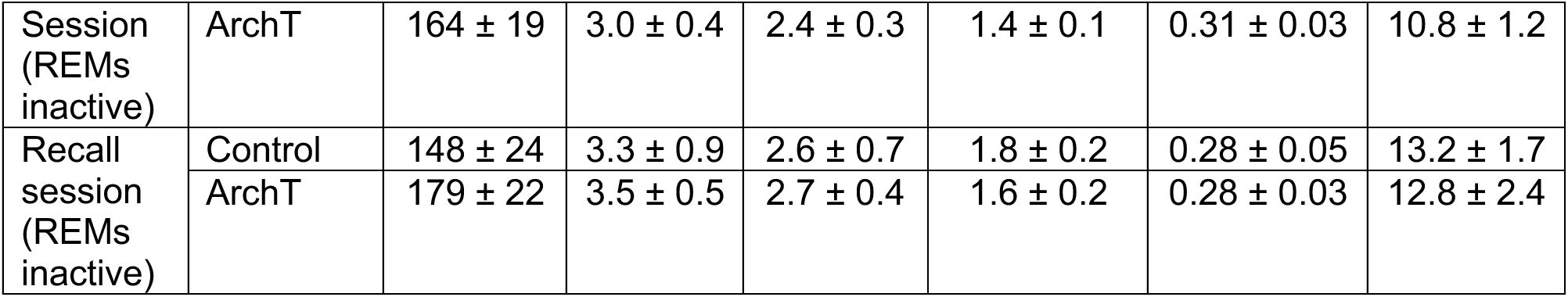
Analysis of REMs-active and REMs-inactive place cell characteristics between Control and ArchT mice during the encoding and recall test sessions. (n=18,19 cells from 7 mice for REMs-active and REMs-inactive place cells, respectively (Control), n=14,27 cells from 6 mice for REMs-active and REM-inactive place cells, respectively (ArchT); non-significance indicated by absence of symbols, unpaired t test (place field size) and Mann Whitney tests (max rate, mean rate, information content, sparsity, selectivity) were used). All values are presented as mean ± SEM.

## Methods

### Data acquisition

#### Subjects

All mice were treated according to protocols and guidelines approved by McGill University and the Canadian Council of Animal Care. VGAT-ires-Cre (VGAT::Cre) transgenic mice (The Jackson Laboratory, stock number 016962) were used in order to selectively target Cre recombinase expression (optogenetic inhibition) to GABAergic neurons. When not being used for experiments, mice were housed at a regulated temperature (22° C) and humidity (30-50%), were kept on a regular circadian cycle (12 h: 12 h light-dark cycle, lights on at 7:30 PM), and had access to food and water *ad libitum* unless otherwise stated.

#### Virus-mediated targeting of archaerhodopsin expression to GABAergic neurons in the medial septum

At ∼10 weeks age, male VGAT::Cre mice were anesthetized with isoflurane (5% induction, 1-2% maintenance) and subsequently placed in a stereotaxic frame. The head of the mouse was then adjusted such that the Lamda and Bregma were in the same horizontal plane. A hole was then drilled in the skull lateral to the midline (AP, +0.86; ML, -0.5; all coordinates relative to Bregma). 0.6 µL of AAVdj-Ef1α-DIO-ArchT-eYFP (plasmids were provided by Dr. K. Deisseroth, viral vectors were packaged at Vollum Vector Core, University of Washington, Washington, USA) was then injected into the medial septum (MS) (anteroposterior (AP), +0.86; mediolateral (ML), 0.0; dorsoventral (DV), -4.5) through a 28 G cannula (Plastics One, Inc., Roanoke, VA, USA) that was angled at ∼6.4 ° (in the ML axis). An injection rate of 50 ηL/min was used in order to minimize disruption of brain tissue at the injection site. Mice were given at least 1 month of rest following the injection to ensure full recovery from the surgery and allow for construct expression in GABAergic neurons within the medial septum (MSGABA).

#### Microdrive and optic fiber implantation for in vivo experiments

∼4 weeks after virus injection, mice were anesthetized with isoflurane (5% induction, 1-2% maintenance). Mice were then placed in a stereotaxic frame and the head was adjusted such that the Lambda and Bregma were in the same horizontal plane. To facilitate single-unit recordings in area CA1 of the hippocampus, a ∼1.5 mm diameter circular potion of the skull overlying the dorsal hippocampus in the right hemisphere (center coordinates: anteroposterior (AP), -1.9; mediolateral (ML), 1.5) was first removed, followed by the dura below. A microdrive (Axona, Inc.) containing 4 independently movable gold-plated tetrodes (150-250 kΩ impedance at 1000 Hz prior to implant) was then placed on the surface of the exposed brain and the base was covered with silicon polymer (Kwik-Sil, World Precision Instruments). The microdrive was then secured in place using 3 anchor screws and dental cement. A wire positioned above the cerebellum served as ground, and an electromyogram (EMG) electrode consisting of stranded tungsten wires (Medwire) was also inserted into the nuchal muscle to record postural tone. To enable light delivery to the transfection zone within the MS, a hole was then drilled in the skull above and lateral to the medial septum (AP, +0.86; ML, -0.5). An optic fiber implant, constructed by cementing a cleaved piece of optic fiber ∼ 12 mm in length (Thorlabs, Inc.) into a ceramic ferrule (Precision Fiber Products, Inc., Milpitas, CA, USA), was then lowered through the hole at an angle of ∼4.5 ° (in the ML axis) until the end of the optic fiber was located just above the medial septum (AP, +0.86; ML, -0.2; DV, -3.83). The entire implant was then secured with further dental cement and tetrodes were subsequently lowered 0.5 mm. Once the dental cement was dry, black nail polish was applied to the surface of the entire implant to prevent excessive disruption of mice due to light emanating from the implant during laser-induced optogenetic inhibition in later experiments. Mice were then returned to their home cage where they were allowed to rest for at least 1 week in order to recover from the surgery.

#### Post-surgery habituation to recording setup

Mice were allowed to recover undisturbed for at least 1 week following implantation of the microdrive and optic fiber. Once recovered, mice were water-deprived (1 mL / day) for the duration of subsequent experiments. Daily checks of weight were used to monitor the water-deprivation procedure; if the weight dropped below 85 % of their *ad libitum* value, additional water was provided until the weight exceeded this threshold. In parallel with the water deprivation protocol, mice also began the procedure for habituation to the general recording setup. This involved daily sessions ∼30-60 minutes in duration in which mice were transferred to a separate ‘habituation’ room within the behavioral testing laboratory and allowed to freely explore a 75 x 75 cm open field. A head stage pre-amplifier tether (Neuralynx, Inc., Boseman, MT, USA) was attached to the connector at the top of the microdrive throughout each session to ensure full habituation to the general recording environment that would be used for subsequent experiments. To encourage the development of foraging behavior in the test area by water-deprived mice, single water drops were scattered evenly throughout the open field.

#### Optimization of tetrode locations in CA1

The optimization of tetrode locations to the cell layer of CA1 was performed simultaneously with the habituation procedure discussed above. Ripple activity, which is maximal near the CA1 cell layer, was monitored from local field potential (LFP) recordings obtained from tetrodes each day during periods of the habituation session when mice were sleeping. Each tetrode was slowly advanced towards the CA1 cell layer at a rate of 25 µm per day until there was a noticeable decline in day-to-day ripple activity. Ripple activity was subsequently monitored over several days to ensure stability, at which point tetrodes were no longer turned. Cumulatively, this procedure took ∼4 weeks to complete.

#### Novel object place recognition procedure

Following habituation and completion of the tetrode location optimization procedure, behavioral experiments began. To enable analysis of the effect of disrupting REMs MSGABA activity on spatial memory performance, we used a novel object place recognition test ^14,16^. Experiments were initially organized into 3 groups (conditions), called ‘ArchT’, ‘ArchT control’, and ‘ArchT REM control’, and each mouse contributed experimental data for the ArchT group as well as 1 of the control groups. A single mouse did not contribute experimental data for more than 1 group, and the type and order of conditions being tested were both determined at random. Only 1 experiment occurred per day. Regardless of the condition being tested the test area consisted of an open field measuring 60 x 60 cm. The 4 walls surrounding the open field measured 30 cm in height and included distinct markings on each to aid spatial orientation. The test procedure always commenced immediately before the start of the light cycle (7:30 AM) in order to maximize sleep activity during the post-encoding rest session. The day of an experiment, the test area was first prepared in a recording room within the behavioral testing laboratory that the mouse had never been in previously. A pair of identical objects (∼5 cm in diameter and ∼200 g in weight), called ‘Object 1’ and ‘Object 2’, were then placed in randomly assigned quadrants within the open field with each having the centroid being 10 cm away from either wall forming a given corner. All key features of the test area, including room used (option of 2 different recording rooms within the behavioral testing laboratory), test area wall color (2 options: all 4 walls black or all 4 walls white) and distinct markings (contrasting wall colors), and types of identical objects (option of 2 pairs of identical objects, each pair of objects is differently shaped yet has a similar size and weight to the other) were each determined at random with no feature being used more than once on the same mouse. Once the test area was prepared, the mouse being tested was transferred to the recording room and the head stage pre-amplifier tether was attached to the connector at the top of the microdrive. To enable light delivery to the medial septum, a light weight optic fiber patch cord constructed in house from basic components (Thorlabs, Inc.) was also attached to the top of the fiber optic implant on the mouse. The mouse was then placed into the test area for a spatial memory ‘encoding’ session which lasted 30 minutes. During the first 10 minutes of this period, mice were allowed to explore the open field at will without any interference from the experimenter in order to measure object interaction. A period of 10 minutes was used because mice typically began to lose interest in objects after ∼15 minutes of initial exploration. After 15 minutes had elapsed, water drops were evenly placed at regular intervals throughout the test area to ensure that spike data associated with locations covering the entire open field was obtained; to this end, cumulative coverage of the test area by the mouse was virtually traced (tracer path width = 1 cm) and continuously updated in real time using Cheetah Software (Neuralynx, Inc., Boseman, MT, USA) in order to ensure that mice covered the entire test area fully at minimum 2 times. At the conclusion of the encoding session, mice were removed from the test area and transferred to the habituation room for a 4 h post-encoding ‘rest’ period. During this session, mice were kept in an open top Plexiglas cage with wood chip bedding and left undisturbed. LFP and EMG signals were continuously monitored by an experimenter. Whenever mice in the ArchT condition entered into REMs, orange light (594 nm wavelength) was continuously delivered to the MS from a laser (Laserglow Technologies) via the optic fiber patch cord until the mouse transitioned out of REMs. Mice in the ArchT control group did not have light delivered to the MS at any point during behavioral testing. For mice in the ArchT REM control group, the duration of each REMs episode was noted; however, in contrast to the ArchT group, delivery of laser light to the MS did not begin immediately after detection of REMs, but was instead delayed by ∼5 minutes from the end of the REMs episode. Following this delay laser light was continuously delivered to the MS for a duration equal to that of the preceding REMs episode. In rare cases, a mouse would enter into REMs again during delayed MSGABA inhibition, at which point the laser was turned off. This method thus produced a ‘REMs-like’ pattern of optical MSGABA inhibition that almost entirely avoided REMs. For both conditions involving light-induced silencing of MSGABA neurons (ArchT and ArchT REM control groups), the laser power was adjusted so that an estimated light intensity of 15 mW was emitted from the end of the optic fiber implant located in the MS. This light intensity was previously found to be highly effective in silencing MSGABA neurons without inducing tissue damage or having other unintended consequences on sleeping behavior ^14^. Following completion of the 4 h post-encoding rest period, mice were returned to the open field test area that had been used for the encoding session 4 h prior in order to complete the ‘recall’ test session. The protocol and setup of the test area were identical to those used for the earlier encoding session, with the lone exception being that one of the objects (Object 2) had been moved to a new randomly assigned quadrant. Following completion of the recall session, mice were returned to their home cage in the animal housing facility. The open field test area and objects were thoroughly cleaned with soap and water immediately before and after both the encoding and recall sessions to remove any unintentional odor cues. Because we previously established that the ArchT control and ArchT REM control conditions did not produce differential results ^14^ and our initial experiments for the current study also supported this conclusion, we opted to combine the data available for the two control conditions in the present study (n=5, 2 experiments for the ArchT control and ArchT REM control conditions, respectively) into a single control group (‘ArchT Control’) in order to minimize the number of mice used and simplify the overall experimental design.

#### Electrophysiological data acquisition

LFP, and EMG data were recorded throughout the encoding, rest, and recall sessions. All recorded signals were first amplified by the head stage pre-amplifier tether before being digitized at 32 000 Hz using a Digital Lynx SX recording system and Cheetah Software and saved to a hard disk. A TTL connection between the laser and Digital Lynx SX recording system was used to precisely time stamp laser activity to electrophysiology data obtained during the rest period. For acquisition of spike data, spike waveforms with above threshold amplitudes (>65 µV) were automatically detected in recordings from constituent electrodes of each tetrode during encoding, rest, and recall sessions. Waveforms associated with detected above threshold spikes were subsequently saved across all electrode channels comprising a given tetrode using Cheetah Software.

#### Video-tracking of position during encoding and recall sessions

The position of mice during the encoding and recall sessions was recorded at 30 Hz (730 x 480 pixels) using a camera purchased from Neuralynx (Neuralynx, Inc., Boseman, MT, USA). The camera was positioned directly above the center of the open field test area and the distance (cm) per pixel, the location of all four corners of the square test area, and object locations were all calibrated prior to each recording session to help facilitate subsequent data processing and analysis steps. The location of the animal at each frame was automatically estimated by Cheetah Software during the experimental sessions as the midpoint between two green and red diodes located on either side of the head stage pre-amplifier. This data was represented by time stamped X- and Y-coordinates relative to the camera field of view which were subsequently saved, along with raw video data, to a hard disk.

### Data analysis

Unless otherwise stated, all data analysis was performed using custom written scripts in Matlab (The MathWorks, Inc.).

#### Novel object place recognition analysis

For assessment of novel object place recognition memory, the amount of time mice spent investigating Object 1 and Object 2 was determined for both the encoding and recall sessions. This was accomplished by displaying video data in Matlab and performing a controllable frame-by-frame playback. For all test sessions, frames where the mouse was directly investigating Object 1 or Object 2 were manually labelled. Direct investigation was defined as there being less than 0.5 cm of visible space between the object and the nose of the mouse during a given frame. Periodically, mice would climb on top of an object and survey the test area; frames associated with these time points were not counted as object investigation despite the prior criteria typically being satisfied in this case. Following completion of object interaction analysis, the Object 2 discrimination index was calculated for the first 10 minutes of both the encoding and recall sessions using the following formula (equation 1):

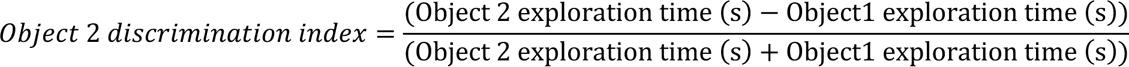

Since mice preferentially investigate novelty, the discrimination index for Object 2, whose position was changed in the recall session relative to encoding session, is considered as a metric of spatial memory; increasingly positive values during the recall session indicate that a mouse has intact memory of the spatial configuration during the prior encoding session.

#### Movement analysis during novel object place recognition encoding and recall sessions

Mouse movement during the encoding and recall sessions was assessed through analysis performed on X- and Y-coordinate data automatically calculated at 30 Hz sampling rate during each session. Raw data values were first imported into Matlab and data time points corresponding to test periods were isolated. The isolated coordinate data was then plotted and manually scanned to control for the presence of erroneous tracking data points due to the reflection of light emitted from the LEDs off of the walls enclosing the test area. Such errors were easily identified as points occurring outside of the calibrated bounds of the test area and were almost always singe isolated events, although in rare cases as many as 3 consecutive frames were affected by inaccurate tracking. In all cases, the errors were easily corrected by automatically interpolating the intermediate coordinate(s) between the two accurate flanking data values. Corrected 30 Hz tracking data was then saved for later use in the analysis of place cell activity during the encoding and recall sessions (see below). For basic movement analysis, 30 Hz tracking data was first subsampled to 1 Hz frequency to prevent isolated head movements and ‘jitter’ in the tracking data signal from artificially inflating cumulative distance values. The cumulative distance, mean speed, and amount of time spent in each quadrant of the test area was then automatically calculated using the subsampled tracking data and test area corner coordinate calibration values. Only the first 10 minutes of the encoding and recall sessions were analyzed in order to avoid motivational changes due to the subsequent addition of water droplets in the test area from unintentionally influencing the results.

#### Vigilance state architecture analysis

For quantitative and qualitative analysis of sleep and wake activity during the 4 h rest period of experiments, raw EMG and tetrode-derived LFP data files were first imported into Matlab and downsampled to 1000 Hz. The best (largest and most stable) LFP signal was then plotted along with EMG data. The major phases of the sleep-wake cycle occurring throughout the entire rest session were subsequently scored in 5 s epochs using the fast Fourier transform (FFT; ‘fft’ function in Matlab) of the LFP recording in combination with the EMG signal ^14^. Epochs of wakefulness were identified by a binned ‘theta’ (4-10 Hz, ‘θ’) to ‘delta’ (1-4 Hz, ‘δ’) power ratio greater than 1 with periods of high-amplitude movement-associated EMG activity which typically lasted several seconds. Epochs of NREMs were identified by a binned θ / δ power ratio < 1 and a lack of high-amplitude EMG activity reflecting behavioral quiescence. Epochs of REMs followed and preceded NREMs and wakefulness, respectively, and were identified in control experiments by a binned θ / δ power ratio greater than 1 and a completely flat EMG signal with the exception being relatively brief and phasic (< 1 s in duration) periods of high-amplitude EMG activity associated with myoclonic muscle twitches. In ArchT experiments, where theta power during REMs was significantly reduced, sustained muscle atonia in conjunction with behavioral quiescence were highly reliable indicators of ongoing REMs. Epochs during which state transitions occurred (e.g., REMs to wakefulness) were scored as the state that was present during the majority of the epoch. Once scoring was completed, the hypnogram was saved and subsequently used to determine the state proportions and average state durations for each experiment.

#### Analysis of oscillatory activity during different vigilance states in CA1 LFP data

For general analysis of oscillatory activity during different vigilance states, spectral analysis was completed on LFP data in Matlab using inputs from the Chronux signal processing toolbox ^43^ (paramters: window size = 5 s, step size = 5 s, tapers [3 5]). For each experiment, values were calculated using the best (most optimal combination of signal size and stability) LFP channel available from any of the tetrodes located in the CA1 cell layer. For analyses requiring greater temporal resolution (analysis of spectral activity during CA1 population event activations and theta-gamma phase-amplitude coupling analysis), Morlet wavelet analysis of concatenated REMs LFP data was performed in Matlab using the ‘cwt’ function. The absolute values of resultant complex wavelet transforms were then calculated for 10 ms time periods centered around each population event activation. Note that population event activations within 1 s of temporal discontinuities in concatenated data were omitted from analysis consideration. Analysis-specific details (LFP channel used, frequency bands used) can be found in the main text and relevant figure legends.

#### Detection of NREMs ripple activity in CA1 LFP data

The detection of transient ripple activity during NREMs in CA1 LFP data was completed using a protocol similar to one reported previously ^14,44^. The entire LFP trace was first band-pass filtered (100-250 Hz) using the ‘filtfilt’ function in Matlab and subsequently rectified. Non-overlapping windows of NREMs 30 s in duration were then identified within the rest period, and the following protocol was completed for each. The 30 s window was first divided into 10 ms segments, and the sum of the filtered rectified LFP signal was calculated for all segments. The mean and standard deviation were then found and used to determine the threshold for ripple detection within a given window (mean + 3 standard deviations above the mean). Ripple locations were identified as above threshold locations within the window, and further refined by combining ripples which were separated by less than 30 ms. For analysis of general ripple features (Table S1), data values were calculated using the best (most optimal combination of signal size and stability) LFP channel from any tetrode. For spike-ripple coincidence analysis (Figure 3C), ripple-coincidence of each spike was calculated based on ripple activity detected using the best LFP channel originating from the same tetrode.

#### Isolation of single neural spike data

Single neural units were identified from spike data using the offline spike sorter SpikeSort 3D (Neuralynx, Inc., Boseman, MT, USA). Principal components extracted from the saved spike waveforms as well as spike waveform amplitude were used to separate (cluster) spikes originating from different neurons recorded using the same tetrode. Only clusters that could be fully isolated with clearly defined boundaries and biologically realistic inter-spike intervals (>1 ms) were included in subsequent analysis. Clusters which had atypical average waveforms (i.e., multiple peaks) were excluded. Given the prolonged experiment duration (∼5 h including the encoding, rest, and recall sessions), care was taken to confirm the absence of significant ‘drift’ in the tetrode location in the CA1 cell layer throughout the full course of an experiment. For this, clustering templates generated for the encoding and recall sessions were checked for near-perfect (greater than ∼80 %) overlap of clusters present in both sessions, and only experiments which satisfied these criteria were included in this study. Once all valid clusters had been identified, the time stamp values corresponding to all constituent spikes were saved for each cluster individually for use in later analysis steps.

#### Analysis of neural spiking activity as a function of oscillation phase

To determine whether isolated single neurons preferentially spiked at specific phases of theta and gamma oscillations during REMs, we first calculated the oscillation phase for different frequency bands from tetrode-derived LFP data obtained during REMs episodes. The calculation of phase preference for spike data originating from each neuron was determined using phase data derived from the best (most optimal combination of signal size and stability) LFP channel of the tetrode from which a single unit was isolated. For analysis of the relationship between population event activity during REMs and oscillation phase, data was obtained from the best LFP channel of the tetrode from which the majority of constituent single units were isolated. However, we do not expect that the choice of electrode significantly influences the results as the key characteristics of LFP recordings obtained from different tetrodes were found to be nearly identical. For the calculation of oscillation phases during REMs, the data was first bandpass filtered (4-10 Hz, 30-60 Hz, or 60-100 Hz) using the ‘filtfilt’ function in Matlab to create ‘band specific’ data. The indices of ‘peaks’ and ‘troughs’ were then found in the filtered data using the ‘findpeaks’ Matlab function on regular and inverted values, respectively. Phase values of 180 and 0 zero were then assigned to the indices corresponding to peaks and troughs, respectively, and intermediate values between peak-trough and trough-peak indices pairs were linearly interpolated.

#### Detection of CA1 population event activity during REMs

Automatic detection of CA1 population event activity during REMs was performed using a windowing approach which was based on detecting, for each possible window location in the REMs data of a given experiment, the number of neurons that were active (fired at least 1 spike) within the window bounds. As a first step, the optimal window size to use was found by calculating the average and maximum coactivity values across all windows of each experiment using window sizes ranging from 1-15 ms (Fig S2B). Specifically, the goal was to find the smallest (highest resolution) window size that would capture the near full extent of population event dynamics, which in this study was considered to be the leveling-off point in graphs plotting average and maximum coactivity values calculated for increasing window sizes. Using this criterion, an optimal window size of ∼10 ms in duration was consistently indicated for experiments performed in all groups. Having identified the optimal window size to detect CA1 population event activity during REMs with in this study, the average and standard deviation of coactivity measures calculated for each 10 ms window was calculated across all REMs data for each experiment separately. These values were then used to determine the experiment-specific threshold, which would subsequently be used to identify windows during which significant CA1 population event activity was present, using the following formula:

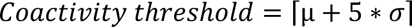

where µ and σ are the average and standard deviation of coactivity measures, respectively, calculated during REMs for a given experiment. The value obtained was then rounded up to the nearest whole number. A relatively high threshold was chosen to minimize the likelihood of false positive CA1 population events being identified at the expense of an increased likelihood of false negative detections. Threshold values calculated for different experiments performed in the same mouse were the same with the exception of 1 mouse; for this experiment, the threshold ultimately used for population event detection during REMs in both was the greatest of the independently calculated values. For each significant CA1 population event detected, the synchrony was measured by determining the minimum interval of time within the corresponding 10 ms window during which the number of active CA1 neurons (neurons firing at least 1 spike) was equal to or greater than the experiment-specific coactivity threshold. For example, if the coactivity threshold calculated for a given experiment was 4 active neurons within a 10 ms period (window), then the synchrony measure for the current population event activation would be the smallest window within which at least 4 neurons fired at least 1 spike.

#### Identification and analysis of place cells

For analysis of spatial tuning of cell activity (place cell analysis) during encoding and recall sessions, only data corresponding to time periods when the speed of the mouse was greater than 2 cm / s was used. Data corresponding to periods when mice were actively investigating either Object 1 or Object 2 was also excluded from place cell analysis. The test area was then divided into 2 cm by 2 cm spatial bins, this bin size being chosen as it was the lowest resolution pixel size which still maintained key occupancy patterns based on comparison to raw occupancy data. The amount of time that a mouse spent in each spatial bin during a given 30-minute test session was then calculated using the 30 Hz tracking data processed in prior analysis steps and subsequently smoothed using a 2-dimensional Gaussian kernel (standard deviation = 4 cm (2 pixels)) using the Matlab ‘smoothdata’ function to produce a smoothed occupancy map. Similarly, a spike map was calculated for each cluster obtained from prior binarized spike raster data by calculating the total number of spikes that occurred in each spatial bin for the given cluster. These raw spike maps were then smoothed with a 2-dimensional Gaussian kernel (standard deviation = 4 cm (2 pixels)), and subsequently divided by the smoothed occupancy map to produce a smoothed rate map. The mean and peak firing rate for each cluster in the test area was then calculated from the smoothed rate maps, and place fields for each cluster were identified by determining the spatial bins in which firing rate was above 60 % of the peak rate. Spatial information content, sparsity, and selectivity were also calculated for spikes in each cluster from the smoothed rate maps using previously established techniques ^45^. The spatial information content of each cluster was calculated using the following formula (equation 2):

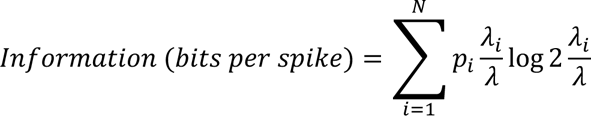

where *i* denotes the current spatial bin, *p*_*i*_ is the occupancy probability in bin *i*, *λ*_*i*_ is the mean spike rate of the current cluster in bin *i*, and *λ* is the overall mean spike rate for the current cluster. The sparsity of spiking activity for a given cluster was calculated using the formula (equation 3):

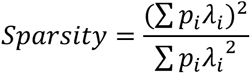

while selectivity was calculated for each cluster by dividing the firing rate of the spatial bin with the maximal value by the mean rate calculated across all spatial bins. The statistical evaluation of spatial tuning of cell activity was performed using a block shuffling procedure for each cluster. Intact tracking data as well as the binarized spike raster constructed using spike time stamp data for a given cluster were split into 30 s blocks and individually reassembled in a random order. The information content of the current shuffled data surrogate was then calculated as described above, and the value stored. This procedure was repeated 100 x; only clusters with intact information content values greater than that calculated for at least 95 corresponding shuffled surrogates were considered as place cells and used for subsequent analysis. The above procedures removed practically all putative interneurons, characterized by narrow (<3 ms) symmetrical waveforms and relatively high firing rates (>10 Hz), from analysis consideration. Only data from clusters which were present and stable in both the encoding and recall sessions were used for comparisons of place cell characteristics between these sessions.

The stability of place cells between the encoding and recall sessions was assessed using 2 methods. The first method calculated the proportion of overlap between place fields of significant place cells during the encoding and recall sessions. The second method calculated the shift in the 2-dimensional place field centroid location from the encoding to the recall session for each significant place cell. Because the ability to accurately detect changes in place field locations (stability) using overlap and centroid measures in encoding versus recall sessions decreases with increasing place field size and is confounded by extreme differences in place field size between the two sessions, we only performed analysis on place cells which fit the following criteria: maximum place field size in either the encoding and recall sessions = 500 cm^2^; maximum difference in place field size between the encoding and recall sessions = 200 cm^2^. These specific values were used as they were found to be highly selective for removal of obvious outlying values.

### Histology

Once all experiments were completed, mice were deeply anesthetized with ketamine/xylazine/acepromazide (100, 16, 3 mg/kg, respectively, intraperitoneal injection). Mice were then perfused intracardially with a PBS solution (1 x PBS-heparine 0.1%, pH 7.4), followed by 4% paraformaldehyde in PBS (PFA). Brains were then extracted and postfixed at 4 ° C overnight before being cryoprotected in 30% sucrose dissolved in PBS for an additional 24 h at 4 ° C. Each brain was then frozen and sectioned at 50 µm using a cryostat. For confirmation of tetrode and optic fiber placement, odd sections were collected, mounted on glass slides, stained with cresyl violet, and coverslipped. For confirmation of archaerhodopsin expression, remaining sections were washed in PBS 1x-Triton (0.3 %) (PBST), incubated in blocking solution (4 % bovine serum albumin (BSA) in PBST) for 60 minutes at room temperature, and subsequently incubated in rabbit anti-GFP (Ab 290, Abcam) diluted 1:5000 in 4 % BSA overnight at 4 ° C. The sections were then incubated in alexa fluor 488 ex anti-rabbit IgG (H+L) (A11008, Invitrogen) diluted 1:1000 in PBST in order to detect the primary antibody for 1 h at room temperature before being mounted on glass slides and coverslipped with Fluoromount-G (0100-01, Southern Biotech). Images of cresyl violet stained sections were obtained using an Axio Observer Carl Zeiss light microscope whereas fluorescent images from immunolabelled sections were collected using an Axio Observer Carl Zeiss fluorescent microscope. Only mice with histologically confirmed electrode and optic fiber placement as well as proper construct expression in the MS were used in the present study.

### Statistics

Unless otherwise stated, all statistical analyses were completed using GraphPad (Dotmatics). *P*<0.05 was considered statistically significant. Experiment-specific test details can be found in the text.

